# z-STED imaging and spectroscopy to investigate nanoscale membrane structure and dynamics

**DOI:** 10.1101/2019.12.28.889923

**Authors:** Aurélien Barbotin, Iztok Urbančič, Silvia Galiani, Christian Eggeling, Martin Booth, Erdinc Sezgin

## Abstract

Super-resolution STED microcopy provides optical resolution beyond the diffraction limit. The resolution can be increased laterally (xy/2D) or axially (z/3D). 2D STED has been extensively used to elucidate the nanoscale membrane structure and dynamics, via imaging or combined with spectroscopy techniques such as fluorescence correlation spectroscopy (FCS) and spectral imaging. On the contrary, z-STED has not been used in this context. Here, we show that a combination of z-STED with FCS or spectral imaging enables us to see previously unobservable aspects of cellular membranes. We show that thanks to an axial resolution of approximately 100 nm, z-STED can be used to distinguish axially close-by membranes, early endocytic vesicles or tubular membrane structures. Combination of z-STED with FCS and spectral imaging showed diffusion dynamics and lipid organization in these structures, respectively.

## Introduction

Cellular membranes are hubs for cellular signalling^1^. They are heterogeneous structures accommodating clusters, domains and nano-assemblies^2^ and this heterogeneity is crucial for cellular signalling^3^. Therefore, there has been extensive effort to resolve the mystery of nanoscale structure and dynamics of cellular membranes. Super-resolution imaging technologies have been extremely useful to shed light on the nanoscale architecture and the supramolecular organisation of the cells^4-10^. Stimulated emission depletion (STED) microscopy is a super-resolution microscopy technique which may be combined with spectroscopic tools to probe the physical and chemical properties of membranes with nanoscale resolution^11-14^. Fluorescence correlation spectroscopy (FCS) is such a spectroscopic tool used to measure molecular mobility in membranes^15, 16^, which is an important parameter to understand the molecular dynamics in cells^15-20^. Combination of STED with FCS (STED-FCS) has been used extensively to address the nanoscale membrane structure^21-27^. Recently STED has also been combined with spectral imaging and polarity-sensitive probes to quantitatively study the nanoscale physiochemical properties of the membrane^11^.

In the context of membrane research, so far only a 2-dimentional depletion scheme (2D STED) increasing the lateral (*xy*) resolution has been used in STED-enhanced spectroscopic measurements. This prevented applications in systems having close-by features along the optical axis (*z*), such as the cellular top and bottom membranes or plasma membrane and internal membranes. A different depletion scheme can be used to increase mostly the axial resolution, relying on a “bottle-shaped” beam for depletion (we will call it z-STED hereafter)^28^. The z-STED depletion pattern has been used both alone and together with 2D STED (3D STED) for imaging^29-35^ and STED-FCS^36-38^ in solution or cytoplasm. However, the exacerbated sensitivity to aberrations^39, 40^ and difficulty of operation of the z-STED depletion pattern have prevented its widespread use. Recently, spatial light modulators (SLMs) have been used in STED microscopy systems to mitigate these challenges, through aberration correction^34, 41-43^ and bespoke calibration protocols^44^.

In this paper, we used SLM-based z-STED with an axial resolution of ≈100 nm to study the nanoscale structure and dynamics of the cell membranes. We show that axially close-by membranes or early endocytic vesicles can be distinguished and studied using z-STED imaging. Moreover, the axial resolution of z-STED allowed extremely sharp optical sectioning which we used to image membrane tubular structures unresolvable by confocal or 2D-STED imaging. Finally, we used a combination of z-STED and FCS to measure diffusion dynamics, and with spectral imaging to assess the lipid organization in close-by membranes.

## Material and Methods

### Bead sample

Microscope slides of 40 nm far-red fluorescent beads were purchased from Abberior Instruments (Germany).

### Preparation of supported lipid bilayers

Supported lipid bilayers (SLBs) were prepared with a spin-coater^45^. The cover slips were cleaned with piranha solution (3:1 sulfuric acid and hydrogen peroxide) beforehand. 1 mg/mL POPC in chloroform/methanol (with 0.01 mol% of Abberior Star Red-labelled phosphatidylethanolamine (PE)) was spin-coated on to a clean coverslip at 3,200 rpm for 30 seconds. The lipid film was rehydrated with SLB buffer (10 mM HEPES and 150 mM NaCl pH 7.4).

### Cells and Maintenance and staining

All cells were maintained at 37 °C and 5% CO_2_. PtK2 Cells were grown in DMEM (Sigma Aldrich) supplemented with 15% FBS (Sigma Aldrich) and 1% L-glutamine (Sigma Aldrich). NIH-3T3 and U2OS were grown in DMEM (Sigma Aldrich) supplemented with 10% FBS (Sigma Aldrich) and 1% L-glutamine (Sigma Aldrich). Red blood cells were obtained from mouse blood. Cells were labelled with the fluorescent lipid Abberior Star Red-PEG-Cholesterol in phenol-red free L15 medium (Sigma Aldrich) at concentration of 0.2 µg/mL for 3-5 min at room temperature. After washing twice with L15, measurements were performed also in L15 medium at room temperature. Each slide was imaged not longer than 30 minutes.

### Optical Setup

We used a custom STED microscope implemented around a commercial RESOLFT microscope from Abberior Instruments (Germany) described in detail in ref^46^, to which we added a spatial light modulator (SLM) (Hamamatsu LCOS X10468-02) in the depletion path, as described in ref^43^. Depletion was ensured by a laser (Spectra-Physics Mai Tai, pulse-stretched by a 40-cm glass rod and a 100-m single-mode fiber) pulsing at a frequency of 80 MHz at a wavelength of 755 nm. STED imaging and FCS were performed using a 640 nm pulsed diode laser (PicoQuant) pulsing at a frequency of 80 MHz. Polarity-sensitive dyes for spectral imaging were excited with a 485 nm pulsed diode laser (PicoQuant). The microscope was equipped with an oil immersion objective lens (Olympus UPLSAPO, 100×/1.4 oil). STED laser power was set to 110 mW, measured in the back focal plane of the objective.

### Alignment of the system

Residual system aberrations in the depletion path of the microscope were removed using the SLM. The depletion beam was imaged by scanning the focus through a sample of scattering gold beads, and the amount of system aberrations present was determined using the sensorless method, using the image standard deviation as image quality metric. Coalignment between the SLM pupil and the objective back-aperture is critical^47^, and we ensured at the beginning of each experiment by inspecting the depletion pattern using scattering gold beads. The detailed SLM-STED calibration protocol we used can be found in ref^44^.

### Image acquisition and processing

Images were acquired with a pixel size of typically 40 nm in the lateral direction and 20 nm in the axial direction and were later resized using python or ImageJ. Pixel dwell times varied between 40 and 160 µs. The contrast of images was enhanced to reduce the visual impact of background contributions due to undepleted side lobes, and to compensate for high intensity variations in images with a large field of view.

### FCS measurements and fitting

The STED microscope was equipped with a hardware correlator from correlator.com (Flex02-08D) operated by Flex software. Abberior Star Red dyes were excited with a 640 nm laser at an excitation power of 2-5 μW. The excitation beam was focussed on membranes by varying its axial position to maximise the signal. The acquisition time of FCS curves was set to 5 seconds. FCS curves were fitted using a custom python script. 2D diffusion model including a triplet state was used to fit the data.

Diffusion coefficients (*D*) were determined from FCS transit times:

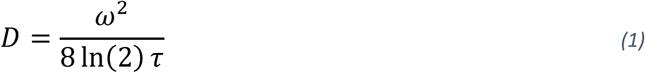

Where *ω* is the full width at half maximum (FWHM) of the Gaussian observation spot and *τ* is the average molecular transit time in the surface area determined with FCS. The FWHM of the confocal spot was determined from images of immobilized fluorescent beads and set to 240 nm. To reliably compare FCS measurements obtained with confocal and STED, we estimated the increase in lateral STED resolution with FCS using supported lipid bilayers (SLBs). Assuming free diffusion, the same diffusion coefficient is expected in both STED (*D*_*s*_) and confocal (*D*_*c*_):

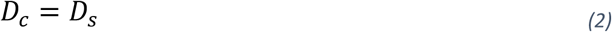

Given equations (1) and (2), the Gaussian lateral FWHM *w*_*s*_ of the STED focus can be estimated as:

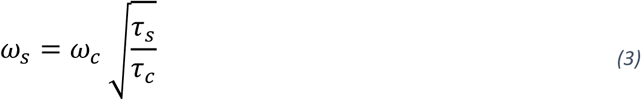

Average number of molecules in the observation area N determined from fitting of individual FCS curves were divided by the measured size of the observation area, to fairly compare confocal and STED recordings. Finally, number of molecules in the observation area were normalised with the confocal value:

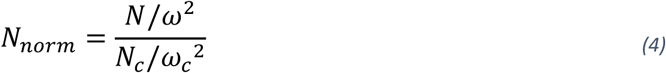

In SLBs, *N*_*c*_ was set to the average value of all confocal recordings. In cells, average number of molecules were normalized separately in each cell to account for variations in label concentrations between different cells.

### Spectral Imaging

We stained the cells with 0.5 µM NR12S in L15 media for 5 minutes at room temperature. Then the cells were washed twice. Imaging was performed in L15 media at room temperature. Each slide was imaged not longer than 30 minutes. We have collected the green channel signal with a 510-590 nm filter (*I*_*G*_) and red channel filter with a 650-730 nm filter (*I*_*R*_). Images were analyzed with the FiJi GP plugin using the equation (5) ^48^.

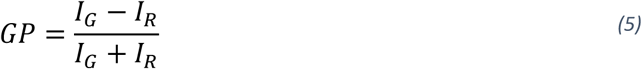

## Results and Discussion

### Axial resolution improvement with z-STED

The most common implementation of STED makes use of a ring-shaped focus (“doughnut”, Figure 1A), created by modulating the phase of the depletion laser with a vortex phase mask. STED imaging with this mask (2D STED) increases the lateral resolution (Figure 1B) but leaves axial resolution unchanged (Figure 1D). In our microscope, the phase mask is created by an SLM, and as such it can be swapped to any other phase mask to change the STED confinement mode without changing the optical layout. Using a top-hat phase mask, a bottle-shaped depletion pattern (Figure 1C) can be created that increases mainly the axial resolution (Figure 1D), but also slightly increases the lateral resolution (Figure 1B). Certain STED microscopes use a combination of the 2D and z-STED depletion patterns to increase the resolution along all dimensions (called 3D STED), at the cost of a higher experimental complexity, lower signal values because of a smaller volume of fluorescence emission, and higher sensitivity to aberrations^49^. In this paper, mainly axial resolution improvement is sought, and therefore z-STED is preferred over 3D-STED.

**Figure 1.**
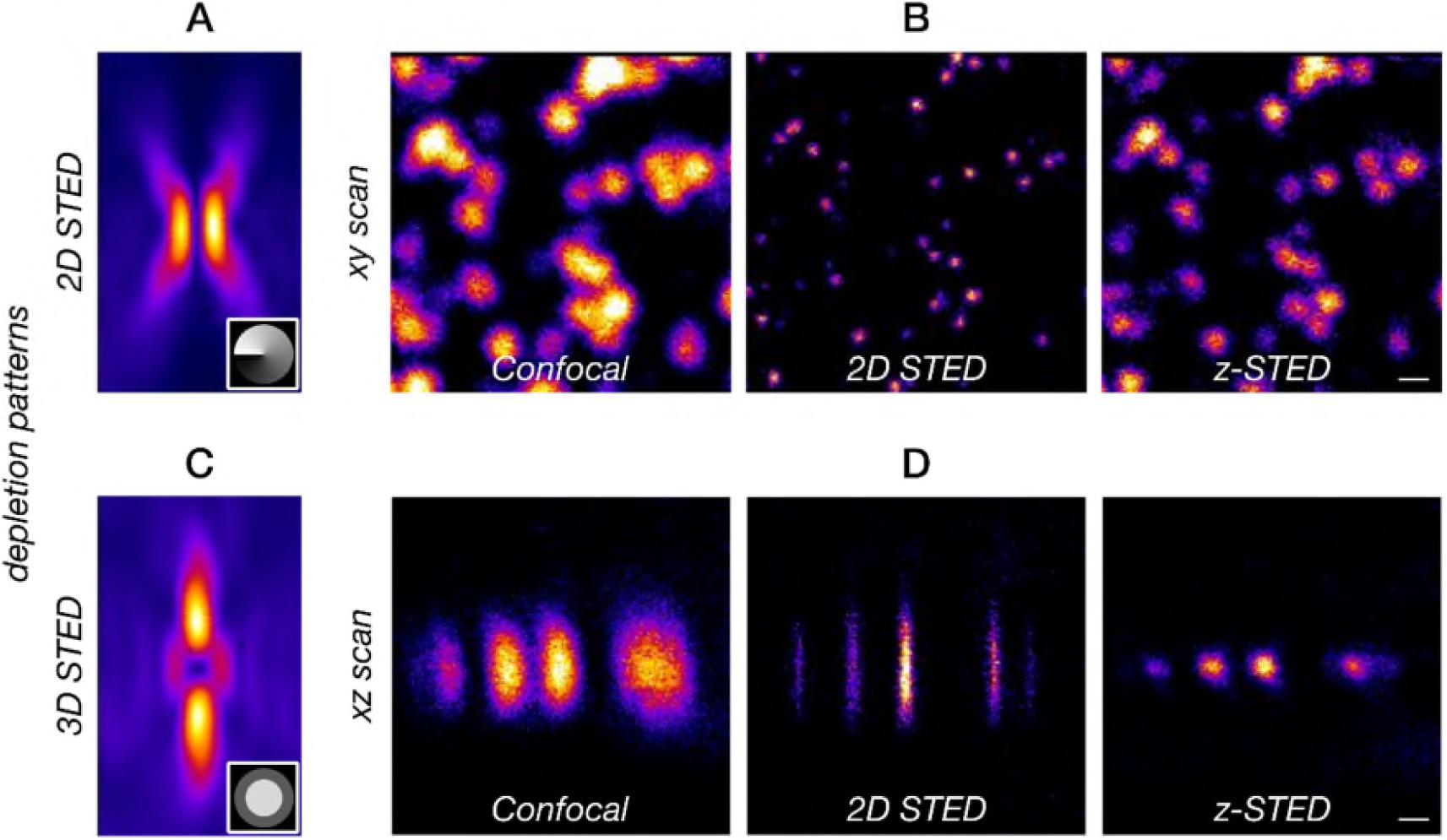
2D and z-STED confinement modes. A and C) Depletion beams of (A) 2D STED and (C) z-STED, visualized by scanning a sample of scattering gold beads through the depletion focus. Insets: phase masks used to create the depletion patterns (grey to black color scale for 0 to 2π phase delay). B and D) xy (B) and xz (D) images of immobilized fluorescent beads, imaged with confocal (left), 2D STED (middle) and z-STED (right). Scale bars are 250 nm.

### Resolving adjacent membranes with z-STED

We first estimated the performance of our z-STED microscope on lipid membranes using a supported lipid bilayer (SLB), which is a single lipid bilayer deposited on a glass surface. This membrane is approximately 5-8 nm thick, which is well below the expected axial resolution of the z-STED microscope, and is therefore an excellent sample to estimate the axial resolution. In confocal imaging, the full-width at half maximum (FWHM) of the axial intensity profile was approximatively 700 nm, while z-STED reduced it to approximatively 100 nm (Figure 2A, B). Undepleted side lobes created a shadow image at approximately 800 nm above and below the membrane, however at a much lower intensity that can be eliminated by image deconvolution (Supplementary Figure S1), or by simply optimizing the contrast of images. In presence of two close-by membranes, such as in Ptk2 cells, the increased axial resolution of z-STED allowed us to resolve cellular top and bottom membranes as close as >100 nm (Figure 2C-E). Similar observations could be made in different cell types such as NIH-3T3, U2OS cells or red blood cells (Supplementary Figure S2). Finally, we used the capability of z-STED to resolve axially close-by membranes to image layered cells (Figure 2F), revealing once again details in images that were inaccessible to confocal images only.

**Figure 2.**
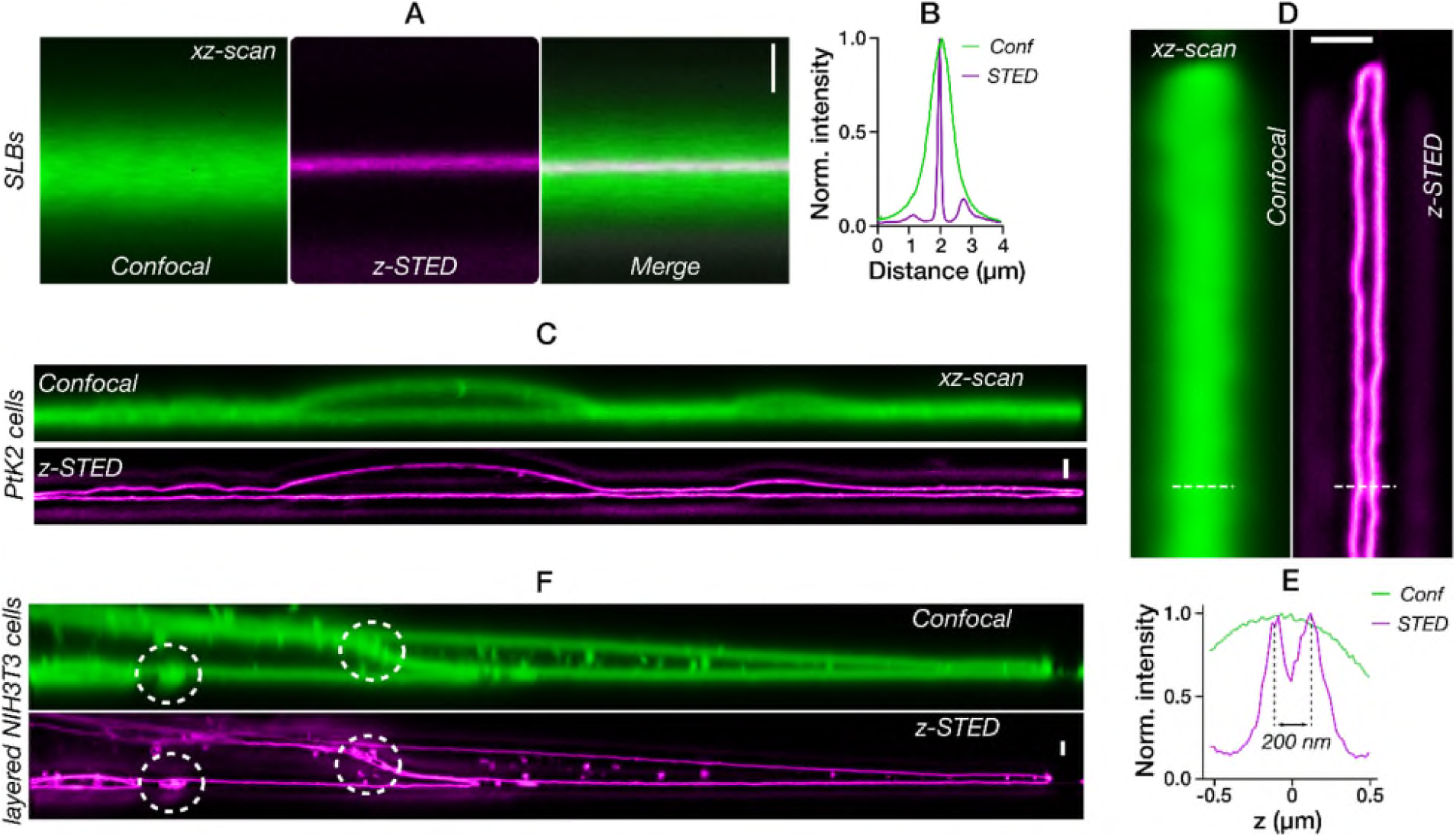
Resolving axially close-by membranes with z-STED. A) Confocal (left), z-STED (middle) and merged (right) images of an SLB, B) Confocal (green) and z-STED (magenta) fluorescence intensity profiles along the axial direction of the SLB shown in A. C) Confocal (green) and z-STED (magenta) images of a live Ptk2 cell. D) Top and bottom membranes of a live Ptk2 cell, imaged with confocal (left) and z-STED (right). E) Intensity profiles along the line drawn in picture D. F) Confocal and z-STED images of layered NIH3T3 cells. z-STED revealed gaps between layers of membranes (white circles). Scale bars are 1 µm and oriented along the optical axis (z direction).

### Resolving submicron structures with z-STED

Submicron structures are common in biology and can have various topologies. For example, submicron tubular structures are crucial for cellular communications^50^. They can usually not be studied with conventional fluorescence microscopes, the resolution of which is insufficient to differentiate, for instance, between stacked and hollow membrane tubes. Such multi-layer membrane patches are common in SLBs, even though it is not clear how these patches can exist despite the hydrophobic repulsion of the lipid acyl chains at the edges of these patches. One possible scenario is the formation of nanotubes at the edges of these patches. We verified this hypothesis using the increased axial resolution provided by z-STED. Imaging the multilayer SLB patches with z-STED revealed tubular structures at the edges (Figure 3A) which could not be resolved in confocal images. We further visualized the networks of smaller patches in SLBs which all turned out to be submicron nanotubes (Figure 3B). The axial (*xz*) image cross sections revealed the tubular nanostructures with great precision. Additionally, this depletion scheme also significantly improved image contrast which was extremely helpful for *xy* images of these structures. When we acquired lateral (*xy*) images and compared the results obtained with confocal, 2D and z-STED (Figure 3C, D), the tubes appeared nearly full in confocal and 2D STED images, while z-STED significantly increased the image contrast by removing out-of-focus light originating from the top and bottom of tubes, allowing the precise visualization of such structures. Similarly, we were able to resolve and improve image resolution and contrast on toroidal structures of red blood cells (RBCs; Figure 3E, F). In both SLBs and RBCs, the quality of 2D STED images was deteriorated by out-of-focus contributions, which further showed the necessity of the excellent axial confinement provided by z-STED. Finally, we showed that z-STED is suitable for imaging early endocytic vesicles, another biologically important structure with a spherical topology (Figure 3G).

**Figure 3.**
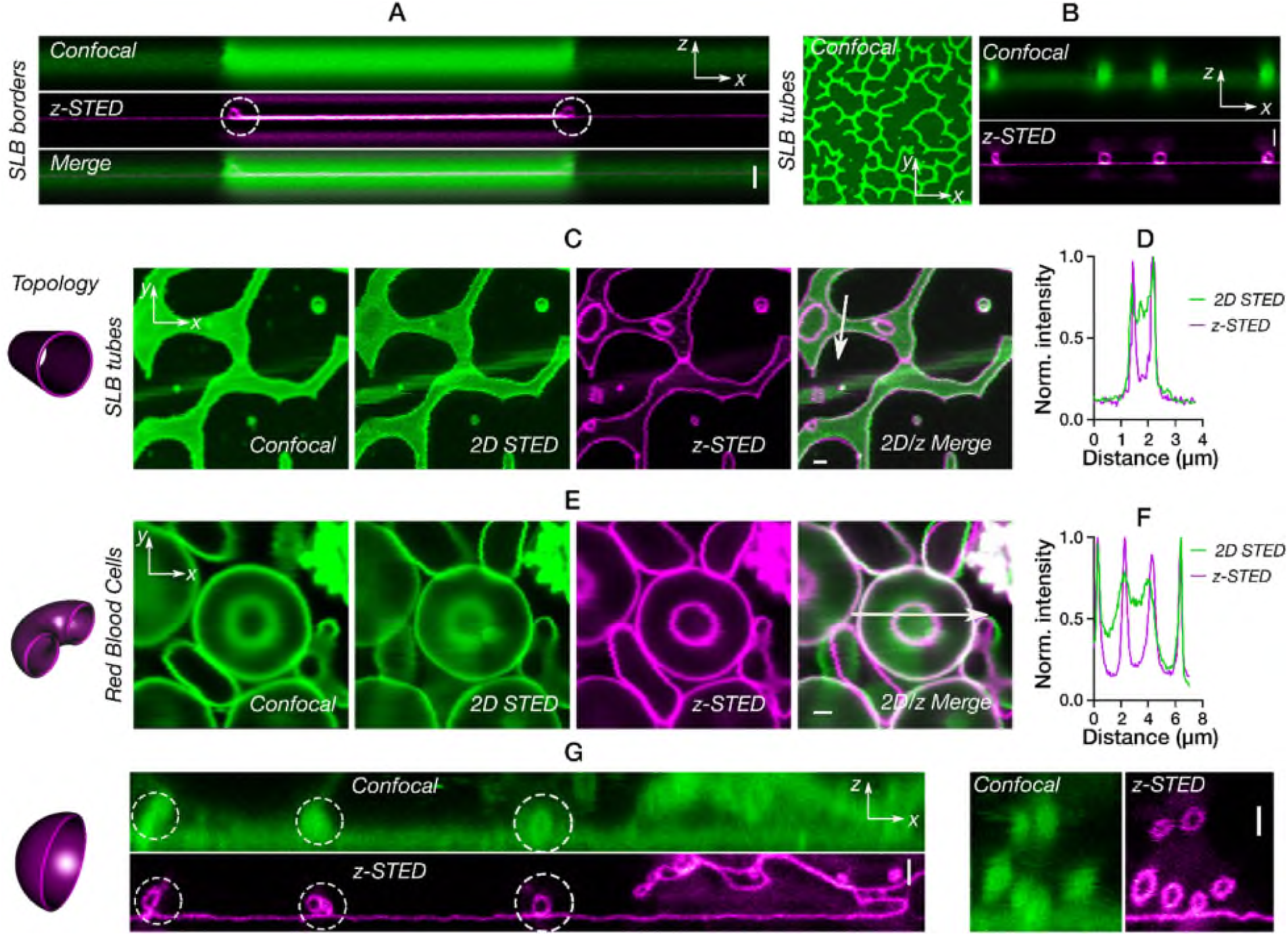
Imaging submicron membrane structures with z-STED. A-B) Tubular structures in SLBs: (A) Confocal (top, green), z-STED (middle, magenta) and merged (bottom) *xz* images of a multilayer structure with tubes at the edges (white circles). B) Lateral (*xy*) confocal (left), axial (*xz*) confocal (top right; green), z-STED (bottom right; magenta) images of hollow networks of multilayered SLB patches. C-D) Membrane tubes in SLBs: (C) *xy* images in confocal, 2D and z-STED modes as labeled with (D) intensity profiles of a tube in 2D and z-STED along the line shown in panel C. E-F) Red blood cells with a toroidal shape: (E) confocal, 2D and z-STED images (*xy* axes) and (F) intensity profiles of a cell in 2D and z-STED along the line shown in panel E. G) Comparison of *xz* cross sections for confocal and z-STED imaging of early endocytic vesicles (white circles). Scalebars are 1 µm. SLBs are labelled with Abberior Star Red-PE, cells are labelled with Abberior Star Red-cholesterol.

### z-STED with fluorescence correlation spectroscopy to investigate membrane dynamics

A unique feature of STED compared to other super-resolution microscopy techniques is the quasi-instantaneous fluorescence switching mechanism employed for reducing the effective observation spot size and thus for increasing the spatial resolution. This speed allows the combination of this technique with spectroscopic techniques such as FCS which requires very high temporal resolution. A few studies applied z-STED together with FCS, yet mainly focusing on cytoplasmic investigations and not on membrane dynamics^36, 37, 43^. Instead, we used here the capability of our z-STED microscope to resolve adjacent membranes (Figure 2, 3) to distinguish their dynamics. We first calibrated our z-STED-FCS measurements on a simple SLB system with confocal and z-STED (Figure 4A, B). With these measurements, we determined the effect of z-STED on diffusion time (τ_D_) and apparent number of molecules in the observation volume (N). We observed an approximately two-fold decrease in values of the lateral transit time τ_D_ through the observation spot for the z-STED recordings (Figure 4C), similar values of the lateral diffusion coefficient D (Figure 4D), and a slightly increased normalized values of the apparent number N of fluorescent molecules in the observation spot (Figure 4E). The decreased values of τ_D_ indicate a 30 % reduction in the lateral size of the observation volume for z-STED (as detailed in Materials and Methods section), which is perfectly in line with the reduction observed for the bead images of Figure 1. Knowing the lateral sizes of the confocal and the z-STED foci, we could calculate the diffusion coefficient of the lipid dye in SLBs (Figure 4D) and compare the average number of molecules per area unit in confocal and z-STED (Figure 4E). We found that the apparent number of molecules per area unit was slightly larger in STED than in confocal (Figure 4E), most likely due to spurious background contributions decreasing the amplitude of the STED curves^36, 37, 51^. Using the calibration of the lateral observation spots obtained with SLBs, we could compare z-STED and confocal FCS measurements in live cells (Figure 4F-I). Particularly, we set out to measure the diffusion speed in two close-by plasma membranes to investigate whether the molecular mobility varies between bottom and top membrane, for instance, because of interactions with the coverslip. Confocal FCS measurements were used to measure the diffusion in the two membranes together, which could not be separated due to limited resolution. We used z-STED-FCS to measure separately the diffusion of molecules in the top and bottom membranes (Figure 4F, G) and did not observe any change in diffusion coefficient *D* between bottom and top membrane or between confocal and STED (Figure 4H). Moreover, there was no difference in normalized average number of molecules between bottom and top membrane. However, we observed that the normalized average number of molecules was twice higher in confocal than with z-STED (Figure 4I), which is consistent with the fact that the confocal FCS measurements were performed on two membranes at once, while only one at a time was measured with z-STED (see also supplementary material for more detailed discussions).

**Figure 4.**
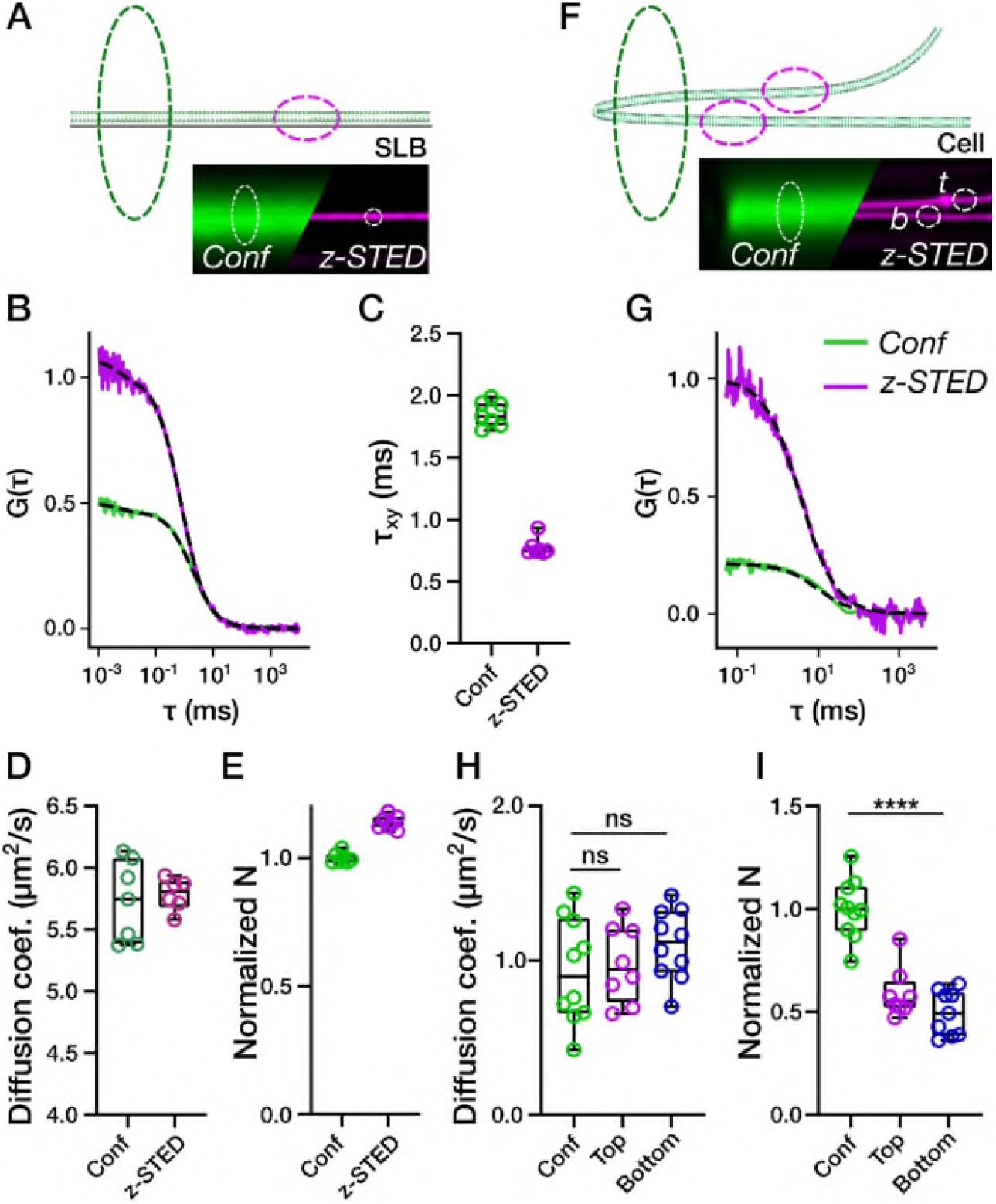
z-STED-FCS measurements in membranes. A-E) FCS measurements in an SLB. (A) Scheme of the experiment: confocal (dotted green ellipsoid) and z-STED-FCS (dotted magenta ellipsoid) measurements were performed on an SLB (lipid bilayer, green). Bottom right: representative confocal and z-STED pictures of an SLB. (B) Representative z-STED and confocal FCS curves on SLB. (C) Average transit times measured in SLBs. D-E) Measured diffusion coefficient (D) and average number of fluorescent molecules per surface area (E) normalized with confocal values measured in SLBs. F-I) FCS measurements in living cells. (F) Scheme of confocal and z-STED-FCS on two close-by membranes (top and bottom) in cells. (G) Representative z-STED and confocal FCS curves in cells. (H) Diffusion coefficient and (I) Molecular density (number of molecules normalized with observation area) in the top and bottom of the cells measured with z-STED-FCS or confocal FCS.

### z-STED combined with spectral imaging to investigate membrane structure

Membrane fluidity is a crucial aspect for membrane bioactivity^52^. An indirect and straightforward way to assess membrane fluidity is the use of polarity sensitive fluorescent probes, whose emission spectra shift with the polarity of the environment. Polarity in membranes generally varies with the hydration level of the bilayer which is itself a function of lipid acyl chain packing. Compared to unsaturated lipids saturated lipids form more tightly packed or ordered membranes where there is less space for water molecules. Recently, we have shown that the custom-synthesized environment sensitive probe NR12S^53^ is accurate for measuring membrane fluidity using 2D STED^11^. We set out to test whether NR12S can also be used with z-STED to characterize and distinguish membrane fluidity in the close-by basal and apical membranes of adherent cells. Live PtK2 cells stained with NR12S were investigated by z-STED by splitting the fluorescence emission into two separate channels: a green (510-590 nm) and red channel (650-730 nm). Fluorescence intensity from more ordered membrane environments increased the relative intensity collected in the green channel, while that from disordered membranes increased the relative intensity in the red channel. Using these fluorescence intensities, general polarization (GP) was calculated as detailed in the methods section as a direct measurement of lipid ordering, with a high GP being associated with a high degree of packing. For the confocal recordings, it was impossible to discern two close-by membranes (Fig. 5A, B), however for z-STED recordings, we could distinguish the top and bottom membranes (Fig. 5A, B) and measure the GP values for each separately (Fig. 5B, C). As expected from the diffusion data (Figure 4), we did not observe any difference between the lipid packing of the top apical and bottom basal membranes of the adherent cells. Along the line, we could also successfully distinguish the plasma membrane from endocytic vesicles and measure their GP separately (Figure 5C) which showed a lower GP for endocytic vesicles compared to plasma membrane. Similarly and as already highlighted in Figure 3, z-STED allows also to increase image contrast in *xy* images due to the reduction in out-of-focus signal, which allows acquisition of high-contrast lateral GP images of structures such as tubes or vesicles in living cells (Supplementary Figure S3).

**Figure 5.**
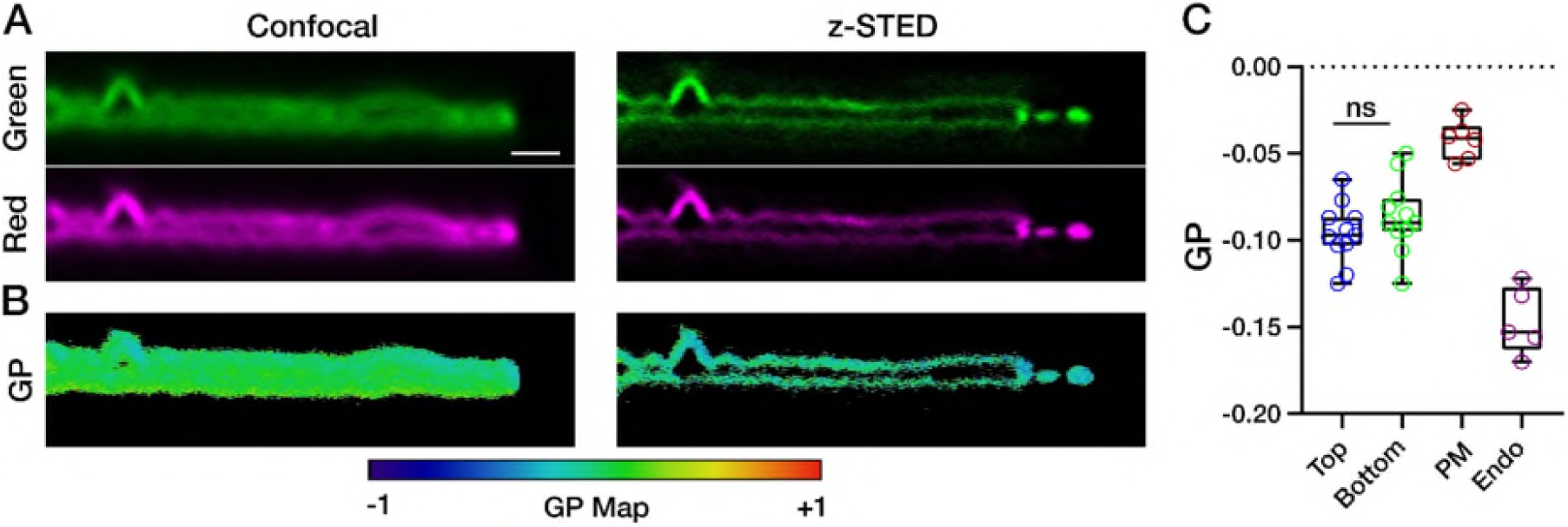
z-STED measurements of lipid order on PtK2 cells using the polarity-sensitive dye NR12S and spectral analysis. A) *xz* images of cells labelled with NR12S, confocal (left) and z-STED (right). B) *xz* confocal and z-STED GP images of the cells shown in panel A, calculated using equation (5), showing two adjacent membranes clearly resolved with z-STED. C) Quantification of GP for bottom vs top membrane of the cells and whole plasma membrane vs early endocytic vesicles. For PM measurements, newly formed endocytic vesicles were avoided. Each data point represents a cell. Scale bar is 1 µm.

## Conclusion

We showed here the efficient use of z-STED for image-and spectroscopy-based studies of the cellular plasma membrane. We observed fine structural details in the z-STED data that could not be resolved in the confocal counterpart. Particularly, we were able to resolve the bottom basal and top apical membranes of various cell types, which are separated by a distance well below the diffraction limited axial resolution. Using this increased axial resolution, we imaged submicron features with different topologies, such as spheres (endocytic vesicles), tubes (in SLB patches) and tori (red blood cells). Finally, we showed that z-STED can be used together with spectroscopic tools such as FCS and spectral imaging coupled with polarity-sensitive dyes. While the former allowed us to study the diffusion in nearby membranes, the latter allowed the observation of minute differences in lipid packing between of plasma membrane and endocytic vesicles.

A limitation of z-STED is the undepleted side lobes caused by an imperfect overlap between the excitation focus and depletion pattern, which created dim intensity shadows above and below continuous structures like membranes. These shadows were however created at a large distance from the focus (approximately 800 nm) and were straightforwardly removed using either background subtraction or image deconvolution. Such side lobes due to undepleted signal were also an issue in other applications, and could successfully be removed by adding a second STED laser pattern to the z-STED^29^ or by engineering the depletion focus to create a better overlap between excitation and depletion, as was previously done in 4pi STED microscopes^54^.

A limitation of STED microscopy is the use of high laser intensities, potentially introducing high phototoxicity and photobleaching, which usually prevent imaging of living cells over long periods of time. This was however less an issue in our current study, since the photobleached fluorescent lipids were vastly replenished due to their fast diffusion. Consequently, we were able to image the same structures across multiple frames without noticeable reduction in intensity as well as cell viability. In this context, the use of diffusing fluorescent lipid analogs offers a robust alternative to other labelling strategies using exchangeable fluorophores^33^.

We could perform FCS measurements in the top and bottom membranes of Ptk2 cells, permitted only by the excellent (≈100 nm) axial resolution provided by z-STED. However, this high precision meant that even small displacements of the membrane along the optical axis significantly biased FCS recordings, effectively reducing the maximum available acquisition times. This problem was solved in 2D using scanning FCS and off-line correction of cellular motion^55^. A different implementation using a deformable mirror for fast z-scanning might also be used to solve this issue in 3D.

## Acknowledgements

We thank Hal Drakesmith Lab for providing red blood cells. We thank the Wolfson Imaging Centre Oxford and the Micron Advanced Bioimaging Unit (Wellcome Trust Strategic Award 091911) for providing microscope facility and financial support. We acknowledge funding by European Research Council (AdOMIS 695140), Wolfson Foundation, the Medical Research Council (MRC, grant number MC_UU_12010/unit programmes G0902418 and MC_UU_12025), MRC/BBSRC/EPSRC (grant number MR/K01577X/1), EPSRC/MRC (grant number EP/L016052/1), the Wellcome Trust (grant ref 104924/14/Z/14), the Deutsche Forschungsgemeinschaft (Research unit 1905 “Structure and function of the peroxisomal translocon”), Jena Excellence Cluster “Balance of the Microverse”, Collaborative Research Center 1278 “Polytarget”), Jena Center of Soft Matter, Oxford-internal funds (John Fell Fund and EPA Cephalosporin Fund), European Commission (MSCA IF 707348 (to IU) and Wellcome Institutional Strategic Support Fund (ISSF). ES is funded by the Newton-Katip Celebi Institutional Links grant (352333122).

## Data Availability

The research materials supporting this article can be accessed by contacting the authors.

