## Supplementary material for "z-STED imaging and spectroscopy to investigate nanoscale membrane structure and dynamics": Supplemantary Information

### Supplementary Information

#### Image deconvolution

Shallow side lobes due to undepleted fluorescence signal were particularly visible when imaging membranes with z-STED. A solution to remove them is to use image deconvolution. First, the point spread function (PSF) of the microscope was estimated. From images of SLBs, we estimated the axial intensity profile of the z-STED PSF (see Figure 2C in main text). The lateral intensity profile was assumed to be Gaussian, with a lateral size estimated from FCS recordings in SLBs estimated to be equal to 160 nm (see Methods section). The PSF was then reconstructed as follows:

$$PSF(x, z) = I(z)\exp\left(-4\log(2)x^2/\omega^2\right) \qquad (1)$$

Where *I(z)* is the axial intensity profile and $\omega$ the lateral FWHM of the PSF. Using this PSF (Figure S1A), images were deconvolved employing the Richardson-Lucy algorithm (20 iterations) in Python using the library scikit-image[1]. Deconvolution improved the sharpness of the images and efficiently suppressed the side lobe contributions (Figure S1).

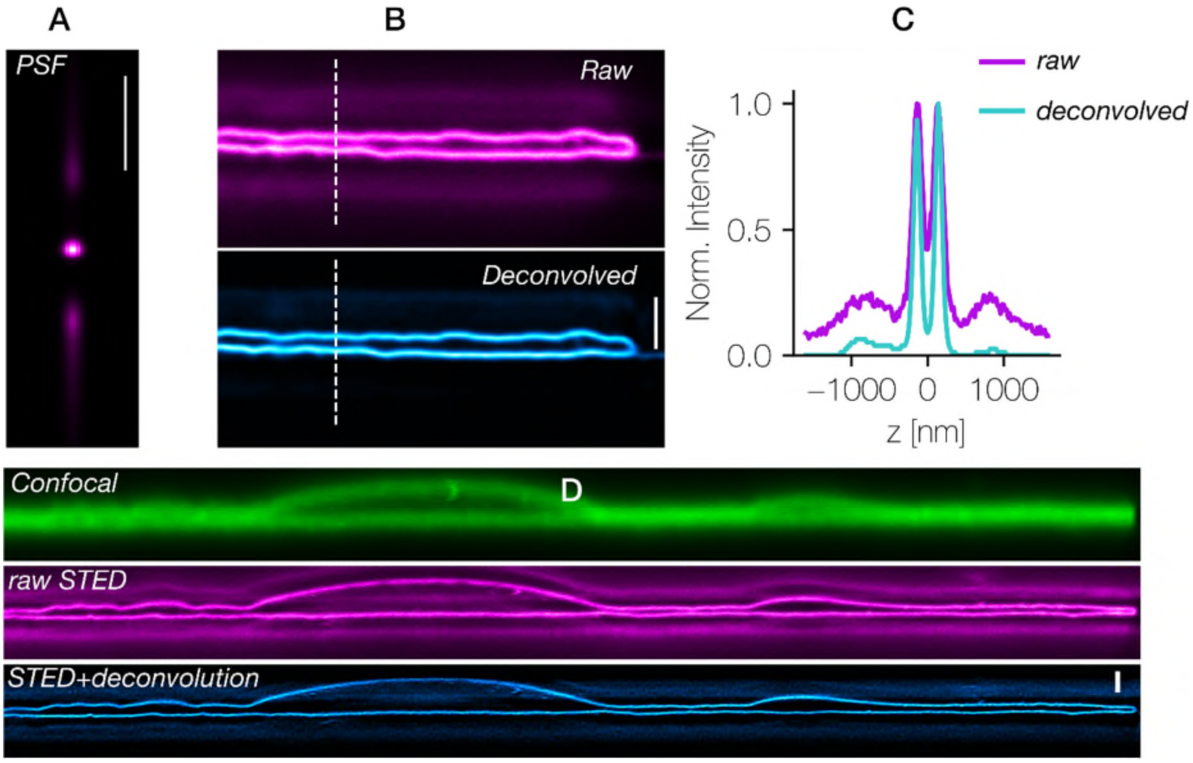


**Figure S1.** Suppressing contribution from side lobes due to undepleted fluorescence signal using image deconvolution. A) PSF of the z-STED microscope, estimated from imaging and FCS of SLBs. B, C) Raw (magenta, top B) and deconvolved (blue, bottom B) of z-STED images of two axially close-by membranes along with intensity profiles (C) along the line marked in B. D) Deconvolution of a z-STED image of a whole Ptk2 cell: comparison of confocal (top, green), raw STED (middle, magenta) and deconvolved z-STED (bottom, cyan). Scale bars 1 µm.

**z-STED imaging of fluorescent lipids in different cell types**

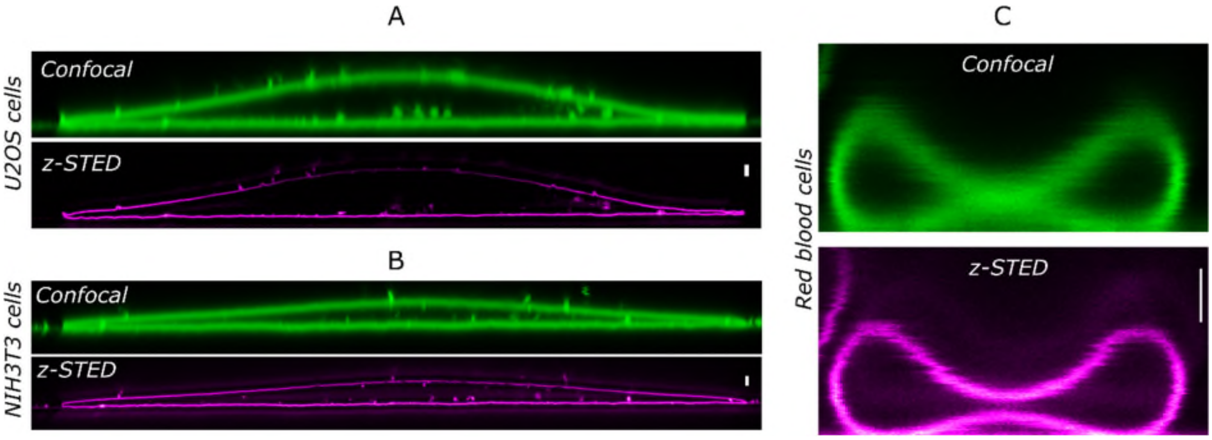


**Figure S2.** Imaging different cell types with z-STED. *xz* images of live A) U2OS, B) NIH3T3, and C) red blood cells, labelled with Abberior Star Red-PEG-Cholesterol. Scale bars 1 µm.

**_xy_ improvement of GP imaging**

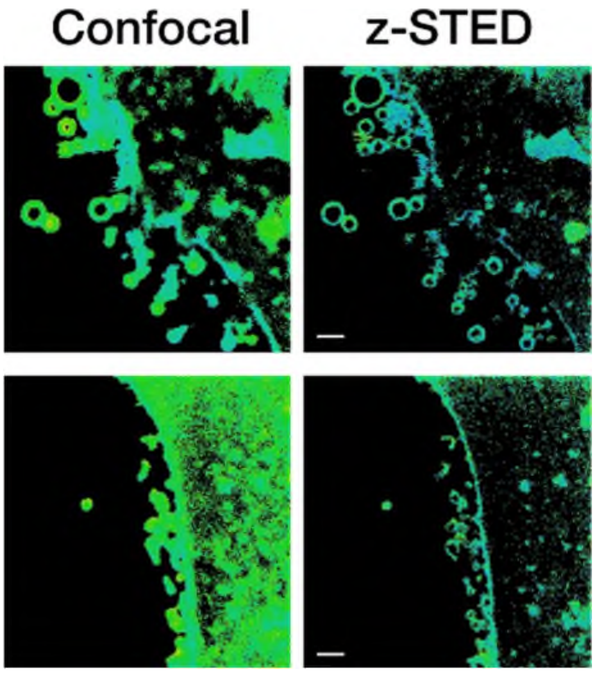


**Figure S3.** *xy* confocal and STED GP images of cells labelled with NR12S probe, showing how increase in contrast in axial direction helps visualizing and characterizing the small vesicles that cannot be resolved with confocal. Scale bars are 1 µm.

**Theoretical analysis of FCS experiments in two close-by membranes**

We compared the results obtained when doing FCS in the top and bottom membranes with z-STED as well as on both at the same time with confocal FCS. It is possible to analytically predict the outcomes of such measurements, if they satisfy three assumptions:

1. The intensity fluctuations in the top and bottom membranes are statistically independent.
2. The top and bottom membranes are close enough so that the confocal cross section is identical at both positions.
3. The label concentration and diffusion speed of the top and bottom membranes are strictly identical.

Following these assumptions, the time-dependent intensity recorded in confocal mode by the detector can be written as:

$$I(t) = I_t(t) + I_b(t) \tag{2}$$

Where *I(t)* is the time-dependent intensity recorded by the detector, $I_t(t)$ is the intensity emitted at the top membrane and $I_b(t)$ the intensity emitted at the bottom membrane.

The autocorrelation function of the time trace fluctuations writes as:

$$G(\tau) = \frac{\langle \delta I(t) \delta I(t+\tau) \rangle}{\langle I(t) \rangle^2} \tag{3}$$

where $\langle\,.\,\rangle$ denotes the time-averaging operator, and $\delta I(t) = I(t) - \langle I(t) \rangle$ represents the temporal intensity fluctuations. Replacing equation 2 in equation 3 and expanding yields:

$$G(\tau) = \frac{\langle \delta I_t(t) \delta I_t(t+\tau) \rangle + \langle \delta I_b(t) \delta I_b(t+\tau) \rangle + \langle \delta I_t(t) \delta I_b(t+\tau) \rangle + \langle \delta I_b(t) \delta I_t(t+\tau) \rangle}{\langle I_t(t) \rangle^2 + \langle I_b(t) \rangle^2 + 2\langle I_b(t) \rangle \langle I_t(t) \rangle} \tag{4}$$

This expression can be greatly simplified using our above assumption. Independence of the fluctuations in the top and bottom membranes (assumption 1) involves that $\langle \delta I_t(t) \delta I_b(t+\tau) \rangle = \langle \delta I_b(t) \delta I_t(t+\tau) \rangle = 0$. Assumptions 2 and 3 state that the average intensities and intensities fluctuations are identical in top and bottom membranes and we can conclude that $\langle I_t(t)^2 \rangle = \langle I_b(t)^2 \rangle$ and $\langle \delta I_t(t) \delta I_t(t+\tau) \rangle = \langle \delta I_b(t) \delta I_b(t+\tau) \rangle$. As such, equation 4 can be simplified as:

$$G(\tau) = \frac{2\langle \delta I_t(t) \delta I_t(t+\tau) \rangle}{4\langle I_t(t) \rangle^2} = \frac{1}{2} G_t(\tau) \tag{5}$$

Where $G_t(\tau)$ is the autocorrelation function obtained if only the top (or bottom) membrane was present. Equation (5) thus means that FCS measurements performed on two axially close-by membranes leaves the transit times (and consequently apparent diffusion coefficient) unchanged, but divides the amplitude by a factor two. This means that the apparent number of fluorescent molecules is twice higher when measuring two membranes at once than when measuring membranes separately. This matches our experimental results (Figure 4, H and I in main text).
